# Socotra Cormorants in the Arabian Gulf represent a large, but isolated population with low genetic diversity

**DOI:** 10.64898/2026.04.01.712451

**Authors:** Noura Mabkhoot Almansoori, Haslina Razali, Sabir B. Muzaffar, Delphine B.H. Chabanne, Ada Natoli, Mohamed Almusallami, Humood Naser, Abdulqader Khamis, Fatimah Al Harthi, Latifa Salem Rashed Aldhaheri, Marwa Mosabeh Bader Alaleeli, Farah M. Al Diwani, Oliver Manlik

## Abstract

The Socotra Cormorant (*Phalacrocorax nigrogularis*) is a threatened seabird endemic to the coastal areas of the Arabian Gulf and the Arabian Sea, two regions separated by the Strait of Hormuz. Conserving threatened species requires clear delineation of population boundaries and the evaluation of genetic diversity. However, information on population structure and genetic variation, necessary for such an assessment, is lacking for the Socotra Cormorants. In this study, we assessed population structure and genetic diversity of Socotra Cormorants using two contrasting genetic markers: (1) maternally inherited mtDNA cytochrome oxidase 1 (COI) and (2) a nuclear non-coding region, β-fibrinogen intron 7 (FIB7). A total of 279 individuals were sampled from four colonies in the Arabian Gulf and one colony on Hasikiyah Island in the Arabian Sea. Findings based on COI-variation suggest that the Arabian Gulf colonies represent one large population with extensive gene flow between Gulf colonies—except for the most distant pair of colonies—but isolated from Hasikiyah in the Arabian Sea. COI-variation indicated significant differentiation between the colonies inside the Gulf and the Hasikiyah colony. This is consistent with the reported distribution patterns, and may reflect phylogeographic processes of the region. The Gulf population showed substantially lower COI-diversity, with significantly lower nucleotide and haplotype diversity compared to Hasikiyah. In contrast, FIB7 results indicated extensive connectivity among colonies, with no detectable population structure or significant differences between the Gulf population and Hasikiyah. This study presents the first characterization of population structure and genetic diversity of Socotra Cormorants. The low genetic diversity coupled with relative isolation of the Gulf Socotra Cormorants raises conservation concerns regarding their long-term viability by potentially reducing fitness and eroding their evolutionary capacity to adapt to environmental change.

**LAY SUMMARY:**

- The Socotra Cormorant is a threatened seabird found in the Arabian Gulf and Arabian Sea, but little was previously known about its population structure and genetic diversity.
- We analyzed 279 birds from five nesting colonies (4 in the Gulf and 1 in the Arabian Sea), using two genetic markers to assess population connectivity and variation.
- We found that the Socotra cormorants inside the Gulf appear to form a large, genetically isolated population with relatively low genetic diversity.
- This is the first study that evaluates population structure and genetic diversity of this endangered seabird.
- This is important information for the conservation of the Gulf Socotra cormorants because low genetic diversity, coupled with relative isolation, is associated with reduced fitness, and suggests that they may have a lower chance to adapt to environmental changes.

## INTRODUCTION

The Socotra Cormorant (*Phalacrocorax nigrogularis*) is a regional endemic seabird found in the Arabian Gulf, the Gulf of Oman, and the southern part of the Arabian peninsula (Khan et al. 2018; BirdLife International 2019). It is classified as ‘Vulnerable’ by the International Union for Conservation of Nature (IUCN) Red List (BirdLife International 2019; IUCN: https://www.iucnredlist.org/species/22696802/155525071). The reasons for this classification include the limited range, habitat loss from coastal development, disturbance at breeding sites, oil pollution, predation and an estimated population decline—from population size estimates of ca. 450,000 −750,000 in 2000 (BirdLife International 2000) to only 220,000 in 2018 (BirdLife International 2019). Most of the global breeding populations, with a total of approximately 110,000 breeding pairs, are inhabiting the Arabian Gulf (86%), and only about 10,000 pairs are found along the coast of Oman in the Arabian Sea on the other side of the Strait of Hormuz (Muzaffar 2015, Muzaffar et al. 2017a; Muzaffar 2020). Currently, the Arabian Gulf hosts about 20 Socotra cormorant colonies (Muzaffar 2020). This excludes several colonies that are no longer used, and individuals that bred on those colonies have presumably moved elsewhere due to human activities, including disturbance on the breeding sites (Jennings 2010, Khan et al. 2018, Muzaffar 2020). The Gulf of Salwah, south of Bahrain and west of Qatar, hosts four colonies, although the current number of colonies is unknown. Most recent estimates from about two decades ago, suggest as many as 25,000 pairs nest on Hawar Island (King 2006), Bahrain, in the Gulf of Salwah. The United Arab Emirates is home to nine colonies, including one on Siniya Island, Umm Al Quwain, which has between 28,000 and 41,000 breeding pairs (Muzaffar 2015, Muzaffar et al. 2017a, Muzaffar et al. 2017b, Whelan et al. 2018). In recent years a new Socotra Cormorant breeding colony was established on the World Islands (W.I.), which is a man-made archipelago off Dubai. The breeding colony on Siniya Island undertakes short-distance directional migration primarily southwest towards islands in Abu Dhabi, although a portion of the population also disperses to the Musandam peninsula (Muzaffar et al. 2017a, 2017b). Individuals breeding on Siniya Island have been observed roosting on the W.I. at the end of the breeding season (Muzaffar et al. 2017a) and it is plausible that Siniya Island breeders could establish alternate roosting and breeding sites, such as the W.I., that are en-route to their non-breeding areas in Abu Dhabi. On rare occasion, Siniya Island birds have migrated over Qatar and into Kuwaiti waters, mingling with birds typically breeding on Hawar Islands, Gulf of Salwa (Muzaffar et al. 2017b; Orben et al. 2025). There is no overlap in distributions between Socotra Cormorants in the Gulf and those of Oman (Arabian Sea), and no record of movements of individuals between the two regions has been observed (Jennings 2010, BirdLife International 2019, Muzaffar 2020). The lack of overlap suggests that the Socotra Cormorants on either side of the Strait of Hormuz may represent two separate populations (Muzaffar 2020). Such population differentiation might be explained by a frontal zone running north along the Strait of Hormuz, which separates plankton, as well as larger faunal assemblages, between the Arabian Gulf and the Arabian Sea (Piontkovski et al. 2019). Such segregation of water masses and biological communities could lead to conditions fostering population differentiation. However, thus far, no study on the Socotra Cormorants has investigated population structure, degree of differentiation and gene flow between colonies in the region. It is unclear whether the colonies inside the Arabian Gulf are isolated from colonies outside the Gulf, or whether the western colony in Bahrain (Hawar Islands) represents a subpopulation to the eastern colonies (e.g. Siniya Island) inside the Gulf.

The delineation of population boundaries and the assessment of genetic diversity is crucial for the management of species of conservation concern (Manel et al. 2003, Moritz 2004, Cross et al. 2016). One powerful approach to reveal population structure and gene flow within and among populations is the dual application of two types of genetic markers (Hoffmann et al. 2009; Manlik et al. 2019a): (1) adaptively neutral (or near neutral) nuclear (nDNA) non-coding regions that allow inference of levels of biparental gene flow among populations; and (2) maternally inherited, non-recombining mitochondrial DNA (mtDNA) sequences, which provide insight into maternal lineages and can reveal patterns of female natal site-fidelity (Birky 2001, Miller-Butterworth et al. 2003, Schlötterer 2004, Manlik et al. 2017). Also, mtDNA variation is commonly used as the basis for assigning evolutionary significant units to guide wildlife management (Moritz 2004). Both mtDNA and nuclear markers have been used to assess fine-scale population structure in a variety of vertebrate populations, including birds (e.g. Cross et al 2016, Fitzgerald et al. 2020, Kimble et al. 2020, Hernández-Soto et al. 2023). For example, population structure and gene flow of Demoiselle Cranes (*Grus virgo*) breeding groups across Europe and Asia were inferred based on one mtDNA locus (control region), indicating recent population fragmentation in Europe (Mudrik et al. 2022). Similarly, phylogeography and Pleistocene refugia of the Little Owl (*Athene noctua*) in Europe was inferred solely from mtDNA variation (cytochrome oxidase I, COI & control region; Pellegrino et al. 2014). Likewise, Blom et al. (2022) used mtDNA loci for population genetic inferences of Bar-tailed Godwits (*Limosa lapponica*) in the Middle East and West Africa. Cryptic lineages, population structure and conservation units of the Eurasian Stone-curlew (*Burhinus oedicnemus*) across Europe, northern Africa and the Middle East were inferred from nDNA microsatellite markers (Baratti et al. (2026).

Genetic diversity is one of the levels of biodiversity for which IUCN recommends protection (McNeely et al. 1990). A certain level of genetic variation is essential for populations to adapt to environmental changes, such as climate change or variation in parasite infection (Frankham 2010, Manlik et al. 2023). Reduction in genetic diversity can lower reproductive output and survival, and ultimately diminishes the evolutionary potential of populations to adapt to environmental changes (Reed and Frankham 2003, Allendorf and Luikart 2008). This close association to fitness and its related demographic consequences, make genetic diversity a crucial variable for the conservation of wildlife populations (DeWoody et al. 2021, Kardos et al. 2021). In a meta-analysis that compared 170 threatened taxa with taxonomically related non-threatened taxa, Spielman et al. (2004) detected lower genetic diversity in 77% of the threatened taxa. In particular, threatened and endangered bird species have significantly lower levels of genetic variation than non-threatened bird species (Li et al. 2016). Although genetic variation cannot identify threat status in the absence of demographic data, predictive accuracy for specific IUCN RedList categories was highest for mtDNA variation in birds (84%), in comparison to other vertebrate taxa and genetic markers, including whole-genome sequences (Schmidt et al. 2023). A case study of the African Sacred Ibis (*Threskiornis aethiopicus*) illustrates the usefulness of mtDNA diversity as a marker for conservation: Ancient mtDNA of *T. aethiopicus* mummies found in the catacombs of Egypt showed substantially greater variation compared to contemporary populations (Wasef et al. 2019, Ku et al. 2023). The aim of this study was to assess population structure and genetic diversity of four Socotra Cormorant colonies inside the Arabian Gulf, and one colony of Oman, Hasikiyah in the Arabian Sea, using two contrasting markers: the maternally inherited mtDNA cytochrome oxidase 1 (COI) and one nuclear, biparentally inherited non-coding marker, the β-fibrinogen intron 7 (FIB7).

## METHODS

### Sample Collection and DNA Extraction

Blood samples of 304 wild Socotra Cormorants were collected in situ between 2019 and 2022 from four nesting sites inside the Arabian Gulf, and one nesting site in the Arabian Sea off the coast of Oman (Figure 1). Within the Arabian Gulf, blood samples from a total of 245 individuals were collected from the following locations: (1) Rubud Al Sharqiya Island (‘HA’; 25°45’4.67“N, 50°47’11.25“E) in the Hawar archipelago of the Kingdom of Bahrain; (2) Butina Island (‘BU’, 24°35’38.26“N, 53° 3’6.16“E, off the coast of western Abu Dhabi (UAE), (3) the World Islands (‘W.I.’, 25°13′00″N, 55°10′00″E), an artificial archipelago off Dubai (UAE; construction start 2003) and (4) Siniya Island (‘SI’, 25°37’3.29“N, 55°37’21.08“E) off Umm Al Quwain. A total of 59 individuals were also sampled from Hasikiya Island (‘HS’, 17°28’31.62“N, 55°36’6.32“E) off the Sultanate of Oman in the Arabian Sea (Figure 1). Note that these sample sites represent nearly the entire distribution of Socotra Cormorants, with the exception of Socotra Island and other possible nesting sites outside the Arabian Gulf (Jennings 2010, BirdLife International 2019, Muzaffar 2020). All samples were collected during the breeding season of Socotra Cormorants in the region (Jennings 2010; Muzaffar et al. 2017b, Muzaffar 2020), between October and November. Relative locations of the sampling sites are shown in the map of Figure 1, along with sample sizes and sampling years for each of the locations.

**FIGURE 1.**
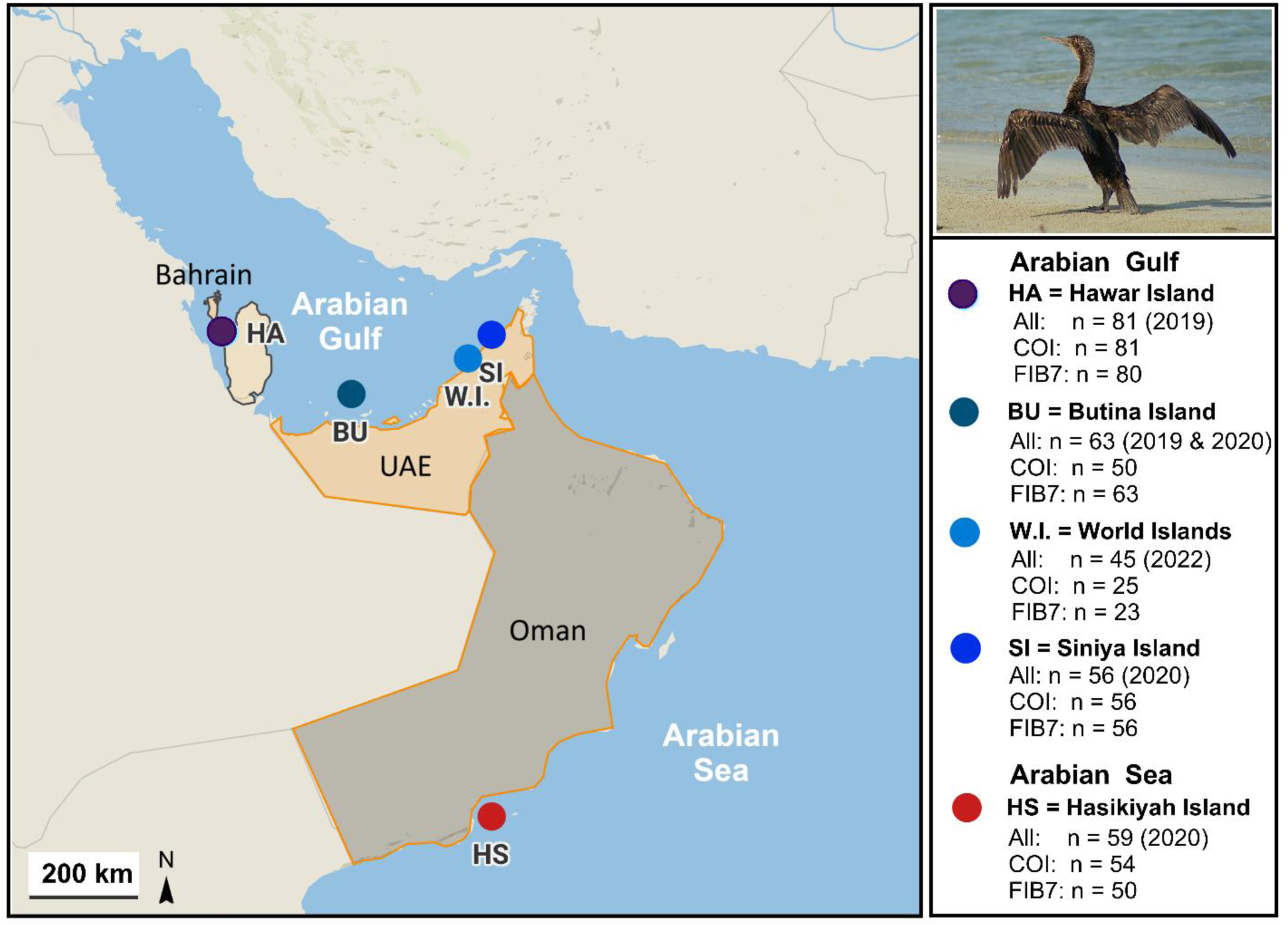
The map shows the sampling locations of Socotra Cormorant breeding sites; inside the Arabian Gulf: HA = Hawar Islands; BU = Butina Island; W.I. = the World Islands; SI = Siniya Island; and in the Arabian Sea: HS = Hasikiyah Island. The sample sizes (‘All: n’) shown in the legend on the right represent the number of individual birds from which blood samples were collected. Sample sizes for analyses, based on mtDNA cytochrome oxidase I (COI) and β-fibrinogen intron 7 (FIB7) are shown below. Numbers in parentheses represent sampling years. Map was generated with Datawrapper (https://app.datawrapper.de/select/map).

We collected blood samples following the general guidelines for avian blood collection by Owen (2011). In brief, upon capturing juvenile (flightless) individuals, blood was drawn from the brachial wing vein, using a 22-25-gauge needle. Common sterile techniques were used to collect the blood, and a cotton ball was applied to the wound to stop the bleeding (Owen 2011). Blood was placed in 100% ethanol immediately upon collection and was then stored at – 20 °C (or – 80 °C for long-term storage) at the United Arab Emirates University (UAEU). Genomic DNA was extracted according to an optimized Qiagen DNeasy Blood & Tissue Kit protocol (Qiagen 2020), followed by ethanol precipitation to desalt and concentrate the DNA. After quantifying the DNA concentration and purity, using the NanoDropTM 2000 Spectrophotometer (Thermo Scientific™ 2009), the DNA working solution was normalized to approximately 20 ng/µL to standardize downstream processes.

### Amplification and Sequencing of Nuclear & Mitochondrial DNA Marker

To assess population differentiation and genetic diversity of the Socotra Cormorants of the region we sequenced two genetic loci: (1) a coding region of mitochondrial DNA, the cytochrome oxidase subunit I gene (COI); (2) one nuclear genetic marker, the β-fibrinogen intron 7 (FIB7). To amplify the COI locus by polymerase chain reaction (PCR), the primer combination FalcoFA (forward: 5’TCAACAAACCACAAAGACATCGGCAC 3’ (Kerr et al. 2007) and COIa (reverse: 5’AGTATAAGCGTCTGGGTAGTC‘3) (Palumbi 1996) were used. This primer pair has been successfully used to amplify and sequence the COI locus in *P. nigrogularis* (Kennedy and Spencer 2014). The primer pair FIB-BI7U (forward: 5’GGAGAAAACAGGACAATGACAATTCAC’3) and FIB-BI7L (reverse 5’TCCCCAGTAGTATCTGCCATTAGGGTT’3) (Prychitko and Moore 1997) was used to amplify FIB7. The FIB7 locus has been amplified and sequenced with this primer pair in other studies on Phalacrocoracidae (Kennedy and Spencer 2014; Kennedy et al. 2019), but not *P. nigrogularis*. PCR mix and thermal cycling conditions for both loci (COI and FIB7) were identical, except for the annealing temperatures. Each PCR mix consisted of 2.5 μL of normalized DNA, 0.5 μL of each primer pair (ca. 0.2 μM), 5 μL 5x FIREPol® Master Mix (including 7.5 mM MgCl_2_) (Solis BioDyne), 0.25 μL of 100× bovine serum albumin (BSA) (to enhance amplification yield), plus ddH_2_O for a total volume of 25 μL. For amplification we used a T100 Thermal Cycler with the following PCR profile: Initial denaturation cycle proceeded at 94°C for 3 min, followed by 40 cycles of denaturation at 94°C for 30 s, annealing at 62.1°C and 50.0°C for COI and FIB7, respectively, for 45 s to 1 min, elongation at 72°C for 1 min, and a final extension at 72°C for 4 min. Note that we tested various annealing temperatures, and 62.1°C and 50.0°C worked best for the amplification of the two loci. ddH_2_O was used as a negative control to rule out the possibility of contamination. Amplicons were visualized by gel electrophoresis on a 1.5% agarose gel (in 1× TBE buffer), stained with RedSafe™ (iNtRON Biotechnology, Inc.). A 1× purple loading dye (New England Biolabs, Inc) was used, and the bands were Documented by the Gel Doc EZ imaging instrument (Bio-Rad). Prior to sequencing, exonuclease shrimp alkaline phosphatase (ExoSAP) was used in a cleanup reaction of the amplicons to remove unincorporated nucleotides. All amplicons were Sanger-sequenced in the forward and reverse direction on an ABI 3730 DNA Analyzer (Applied Biosystems) using BigDye™ Terminator v3.1 sequencing reaction chemistry in the Biology Department Sequencing division at UAEU. More than 10% of the samples were re-amplified and re-sequenced, including all that represented unique haplotypes or SNPs.

### DNA Sequence Alignments & mtDNA Haplotype Network Assessment

The forward and reverse Sanger sequences from a total of 279 individuals (both markers combined; COI: 266; FIB7: 272) were used to form consensus sequences, which were aligned with ClustalW (Thompson et al. 1994) in Geneious Prime® 2022.0.1 (https://www.geneious.com) (Drummond et al. 2011). Sequences with low-quality peaks in the chromatogram were removed. To ascertain the coding sequences of COI, the alignments were translated into amino acid sequences to verify the absence of stop codons and check for silent mutations. We inferred FIB7 haplotypes (alleles) by reconstructing haplotype phases from the unphased sequence alignment data using the coalescent-based Bayesian method Phase (Stephens et al. 2001; Stephens and Donnelly 2003) in DnaSP 6.12.03 (Rozas et al. 2017) with 100 iterations, 1 thinning interval and 100 burn-in iterations. This procedure of constructing haplotypes from biparental unphased sequences has been shown to be reliable across taxa and loci (Stephens and Donnelly 2003; Manlik et al. 2019b). Although COI haplotypes are apparent when inspecting the alignment, DnaSP 6.12.03 was also used to infer COI haplotypes. Subsequently we performed a blastn search to compare inferred COI and FIB7 haplotypes to sequences in the NCBI database. For the mtDNA COI haplotypes, we estimated and visualized the haplotype network in PopART version 1.7 (Leigh and Bryant 2015) using the algorithm for median-joining networks described by Bandelt et al. (1999).

### Assessing Population Structure

To infer population structure, based on the two genetic markers, we took two approaches: (1) we assessed genetic differentiation between each pair of sites via differentiation measures, and (2) we performed a population structure analysis. Differentiation between each pair of sites was assessed by inferring pairwise fixation *F*_ST_ values (Wright 1951, Weir and Cockerham 1984), based on the nuclear marker (FIB7), as well as pairwise *ΦST* values—analogous to standardized *F*_ST_ for haploid data (Meirmans 2006, Meirmans and Hedrick 2011)—based on the mitochondrial DNA marker (COI). Additionally, we estimated Shannon mutual information, *SHua* (Sherwin et al. 2006) indices between each of the five nesting sites. *SHua* is robust over a wide range of population sizes and dispersal rates, and has been reported to better reflect dispersal between sites compared to *F*_ST_ or any other method tested (Sherwin et al. 2006, 2017). We performed an Analysis of Molecular Variance (AMOVA; Excoffier et al. 1992) in GenAlEx 6.5 (Peakall and Smouse 2012) to generate *F*_ST_ and *ΦST* values with associated *p*-values between each pair of sites. GenAlEx 6.5 was also used to estimate pairwise Shannon mutual information (*SHua*) values and their associated *p*-values.

Population structure was analyzed for each genetic marker using the Bayesian model-based clustering method implemented in Structure 2.3.4 (Pritchard et al. 2000). This approach estimates the number of genetic clusters (*K*) in the dataset and assigns individuals to these clusters. Markov Chain Monte Carlo (MCMC) runs were performed for *K* values ranging from one to seven, using a burn-in period of 50,000 iterations followed by 50,000 MCMC repetitions post burn-in, with 25 independent runs per *K* value to ensure consistency in population estimations. This analysis implemented the Locprior model with location priors, as well as the correlated frequency and admixture models, because of the close geographical proximity of some of the sampling sites. To determine the most likely number of clusters (*K*), we applied both the Evanno method (Evanno et al. 2005), which estimates Δ*K* based on the rate of change in the log probability of data [Ln P(D)] across successive *K* values, and the Puechmaille Method (Puechmaille 2016), which uses four alternative estimators: median-based approaches (MedMedK and MaxMedK) and mean-based approaches (MedMeaK and MaxMeaK). These estimators help reduce bias caused by uneven sampling and weak structure. All parameters were estimated using the StructureSelector web interface (Li and Liu 2018), which also generated production of visualizations of population structure as bar plots.

### Assessing and comparing genetic diversity

We assessed and compared a variety of genetic diversity measures based on the COI and FIB7 alignments for each of the colonies as well as for the combined Arabian Gulf population (HA, BU, W.I., SI). The population structure analyses based on mtDNA COI (see Results, Table 1A,B) suggested that the four colonies inside the Arabian Gulf form one well-connected population, separated from HS in the Arabian Sea. We also compared genetic diversity of the Gulf population as a whole to the HS colony in the Arabian Sea (because the prior population structure analyses suggest that the four Gulf colonies represent one population, but separate from HS—see Results). For the purpose of comparing Gulf to HS we used two sampling approaches:

1.) We estimated genetic diversity for the Gulf population, based on all sequences from each of the four Gulf colonies. Specifically, to attain combined genetic diversity measures for the Gulf population, we pooled all sequences from HA, BU, W.I. and SI to generate combined COI and alignments (COI: 212 sequences, 1050 bp; FIB7: 222 sequences, 885 bp).
2.) To address the large differences in sample sizes between the combined Gulf population and HS, as well as to allow for statistical comparisons between the two populations, we generated subsamples of equal sample sizes (i.e., equal number of sequences per subsample) for each of the two populations (Gulf and HS). The ‘Gulf’ subsamples were set up by randomly selecting sequences from each of the four colonies inside the Gulf (HA, BU, SI, W.I.). Likewise, the HS subsamples were generated by randomly selecting an equal number of sequences (subsamples) from HS. Each individual (sequence) was only used once in one of the subsamples.

A) <u>mtDNA COI:</u> For the ‘Gulf’ we set up 23 COI subsamples each containing 9 sequences, and for HS we set up 6 COI subsamples each containing 9 sequences. For the Gulf subsamples, the 9 sequences were randomly drawn from each sampling site (HA, BU, SI, W.I.), with at least one sequence from each of those four sites to represent the diversity of all colonies within the Gulf. To establish subsamples with equal sample sizes, and given that the total number of COI sequence for the Gulf population was 212, we randomly excluded 5 COI sequences to attain the 23 Gulf subsamples containing 9 sequences each (i.e. 207 sequences total). The ‘HS’ mtDNA COI subsamples each contained 9 sequences randomly selected from the pool of HS sequences. For HS, with a total COI sample size of 54 we set up 6 subsamples x 9 sequences (54) for HS. Note that those 54 sequences represented the total COI sample size for HS, so no sequences were excluded from subsampling.
B) <u>FIB7:</u> We generated subsamples of 10 sequences each: 22 subsamples × 10 sequences (220) for the Gulf and 5 subsamples × 10 sequences (50) for HS. Unlike for the COI Gulf subsamples, no sequences were excluded when subsampling the FIB7 sample set, because all the subsamples together represented the total sample size for each population (Gulf: 22 subsamples × 10 sequences = n_Gulf_ = 220; HS: 5 subsamples × 10 sequences = n_HS_ = 50). Each Gulf subsample for FIB7 contained at least one sequence from each of the four colonies (HA, BU, SI, W.I.).

**TABLE 1.** Differentiation measure**s: (A)** Pairwise *ΦST* values based on mitochondrial DNA cytochrome oxidase I (COI) for each pair of nesting sites below the diagonal; associated *p-*values are shown above the diagonal. **(B)** Pairwise *Shua* values based on mitochondrial DNA COI for each pair of nesting sites below the diagonal; associated *p-*values are shown above the diagonal. **(C)** Pairwise *F*_ST_ values based on β-fibrinogen intron 7 (FIB7) for each pair of nesting sites below the diagonal; associated *p-*values are shown above the diagonal. **(D)** Pairwise *Shua* values based on FIB7 for each pair of nesting sites below the diagonal; associated *p-*values are shown above the diagonal. *p*-values that indicate significant differentiation (\**p* < 0.05; \*\**p* ≤ 0.001) are shown in bold. Nesting (sampling) sites are shown on the map on the in the middle: HA = Hawar Island; BU = Butina Island; W.I. = the World Islands; SI = Siniya Island; HS = Hasikiyah Island (Arabian Sea). Map was generated with Datawrapper (https://app.datawrapper.de/select/map).

|  | HA | BU | W.I. | SI | HS |
| --- | --- | --- | --- | --- | --- |
| HA |  | 0.374 | 0.313 | **0.029\*** | **0.001\*\*** |
| BU | 0 |  | 0.332 | 0.064 | **0.001\*\*** |
| W.I. | 0 | 0 |  | 0.133 | **0.001\*\*** |
| SI | 0.036 | 0.036 | 0.026 |  | **0.001\*\*** |
| HS | 0.273 | 0.283 | 0.256 | 0.428 |  |

(B) $Shua$ based on mtDNA COI
|  | HA | BU | W.I. | SI | HS |
| --- | --- | --- | --- | --- | --- |
| HA |  | 0.428 | 0.758 | **0.031\*** | **0.001\*\*** |
| BU | 0.001 |  | 1 | 0.121 | **0.001\*\*** |
| W.I. | 0.001 | 0 |  | 0.334 | **0.001\*\*** |
| SI | 0.004 | 0.003 | 0.002 |  | **0.001\*\*** |
| HS | 0.022 | 0.021 | 0.018 | 0.034 |  |

|  | HA | BU | W.I. | SI | HS |
| --- | --- | --- | --- | --- | --- |
| HA |  | 0.366 | 0.420 | 0.391 | 0.116 |
| BU | 0 |  | 0.358 | 0.397 | 0.127 |
| W.I. | 0 | 0.002 |  | 0.239 | 0.372 |
| SI | 0 | 0 | 0.006 |  | 0.102 |
| HS | 0.008 | 0.008 | 0 | 0.011 |  |

(D) $Shua$ based on FIB7
|  | HA | BU | W.I. | SI | HS |
| --- | --- | --- | --- | --- | --- |
| HA |  | 0.613 | 0.212 | 0.531 | 0.321 |
| BU | 0.003 |  | 0.492 | 0.731 | 0.681 |
| W.I. | 0.006 | 0.007 |  | 0.173 | 0.600 |
| SI | 0.003 | 0.001 | 0.011 |  | 0.167 |
| HS | 0.003 | 0.003 | 0.003 | 0.008 |  |

The following genetic diversity measures were estimated: For both COI and FIB7, we measured the average number of different haplotypes or alleles detected in the alignments (*Na*), the number of private haplotypes (*PH* for COI) or the number of private alleles (*PA* for FIB7), nucleotide diversity (π), as described by Nei (1987; equation 10.5), haplotype diversity (*Hd*; COI) or gene diversity (*Gd*, FIB7) (Nei 1987; equation 8.4), and Shannon’s Index (*I*) (Brown and Weir 1983; Sherwin et al. 2017). Additionally, for the codominant (but non-coding) FIB7 region, we also estimated observed heterozygosity (*H*_O_), unbiased expected heterozygosity (*uHe*; Peakall and Smouse 2012), and Wright’s fixation index (i.e. ‘inbreeding coefficient’, *F*_IS_; Wright 1951) as a measure of heterozygosity excess or deficit in populations for single loci (Hedrick 2009, O’Reilly et al. 2024). We calculated π, *Hd* (for COI)) and *Gd* (for FIB7) using DnaSP 6.12.03 (Rozas et al. 2017). All other diversity measures, namely *Na*, *PH* (COI), *PA* (FIB7), *uh* (COI), *uHE* (FIB7), *H*_O_ (FIB7) and *F*_IS_ (FIB7) were estimated in GenAlEx 6.5 (Peakall and Smouse 2012).

GenAlEx provides estimates for each of these measures per single nucleotide polymorphism (SNP; COI: 10 SNPs, FIB7: 7 SNPs), which allowed us to statistically compare mean values across the SNPs between the colonies, as well as between the combined Gulf population and HS. Specifically, we compared genetic diversity values across the SNPs between the four colonies inside the Arabian Gulf (HA, BU, W.I., SI) using the Kruskal-Wallis tests. This non-parametric test was selected because distributions of values significantly departed from normality and could not be transformed to normal across all sets of comparisons. We also compared genetic diversity measures between the Gulf population and HS across the SNPs. For these pairwise comparisons we used two-tailed Mann-Whitney *U* tests. Likewise, the non-parametric Mann-Whitney *U* test was chosen because values were not normally distributed and could not be transformed to normal using the same transformation for each. Additionally, we compared the mean genetic diversity values derived from the subsampling approach for the Gulf versus HS. Unlike the values based on the entire set of samples, the subsample values for the Gulf and HS were normally distributed, except for the number of different FIB7 alleles (*Na*). Therefore, we used a two-tailed *t*-test to compare those diversity means between Gulf and HS across the subsamples. The comparison of mean values for *Na* across the subsamples between Gulf and HS was performed using a two-tailed Mann-Whitney *U* test. All statistical analyses were performed in GraphPad Prism, version 10.5.0 (www.graphpad.com).

## RESULTS

Of the 304 samples collected, 279 were successfully sequenced, COI: 266, FIB7: 272, while sequencing of 25 samples failed. The final alignment for mtDNA COI, truncated to 1050 bp, included sequences from 266 individuals from all sites (Arabian Gulf: n = 212; Arabian Sea: 54; see Fig 1 for details). Sample sizes for each nesting site for this alignment are reported in Figure 1. The FIB7-alignment, with a truncated sequence length of 885 bp, contained a total of 272 sequences (Arabian Gulf: n = 222; Arabian Sea: n = 50; see Fig 1 for sample sizes for each sampling site). Both markers showed relatively low variability with 10 and 7 single nucleotides (SNPs) for COI (n = 266) and FIB7 (n = 272), respectively, for the two alignments of all individuals from all sampling locations. A total of 12 COI and 7 FIB7 haplotypes were detected in the entire sample set (all four colonies in the Gulf and HS).

### Population differentiation

The haplotype network (Figure 2), depicting the relative proportions and distribution of mtDNA haplotypes in the region, reveals some site-specific haplotype patterns: Haplotype 1 (Hap 1) accounts for about 71% of all haplotypes inside the Arabian Gulf (HA, BU, W.I., SI combined: 151/212), but only about 11% of COI haplotypes in HS (6/54) (Figure 2; Supplementary Material Table S1). Overall, less than 4% of individuals with haplotype 1 (6/157) are from HS, i.e. outside the Arabian Gulf (Figure 2), although HS individuals represent more than 20% of the total sample (54/266). Reversely, haplotype 2 (Hap 2) comprises about 72% (39/54) of all haplotypes of the HS colony, but only about 19% of haplotypes of the Gulf colonies (40/212; Figure 2; Table S1). Accordingly, differentiation between each of the colonies inside the Arabian Gulf (HA, BU, W.I., SI) was relatively low, but high compared to HS in the Arabian Sea (Table 1A,B). Specifically, pairwise *ΦST* and *Shua* values derived from the mtDNA COI sequences were relatively low (*ΦST*: 0.000 to 0.036; *Shua*: 0.000 to 0.004) and non-significant for all pairings within the Arabian Gulf, except for the most distant pairing within Gulf colonies HA-SI, which had relatively low but significant values (*ΦST*: 0.036; *Shua*: 0.004; Table 1A,B). In contrast, pairwise differentiation measures, based on mtDNA COI, were relatively high (*ΦST*: 0.256 to 0.428; *Shua*: 0.018 to 0.034) and highly significant (*p* < 0.001) between each paring of sites inside the Gulf and Hasikiyah (HS) in the Arabian Sea (Table 1A,B). This suggests limited gene flow between the colonies inside the Gulf and HS in the Arabian Sea. Within the Gulf, COI indicated generally low differentiation and high gene flow among colonies, but a small yet significant difference was detected between the two most distant colonies (HA and SI), suggesting subtle matrilineal structuring. In contrast, pairwise *F*_ST_ and *Shua* values inferred from the FIB7 intron were comparatively low for all pairings (*F*_ST_: 0.000 to 0.011; *Shua*: 0.000 to 0.008), and no significant differentiation between any of the nesting sites was detected, based on those values (Table 1C,D).

**FIGURE 2.**
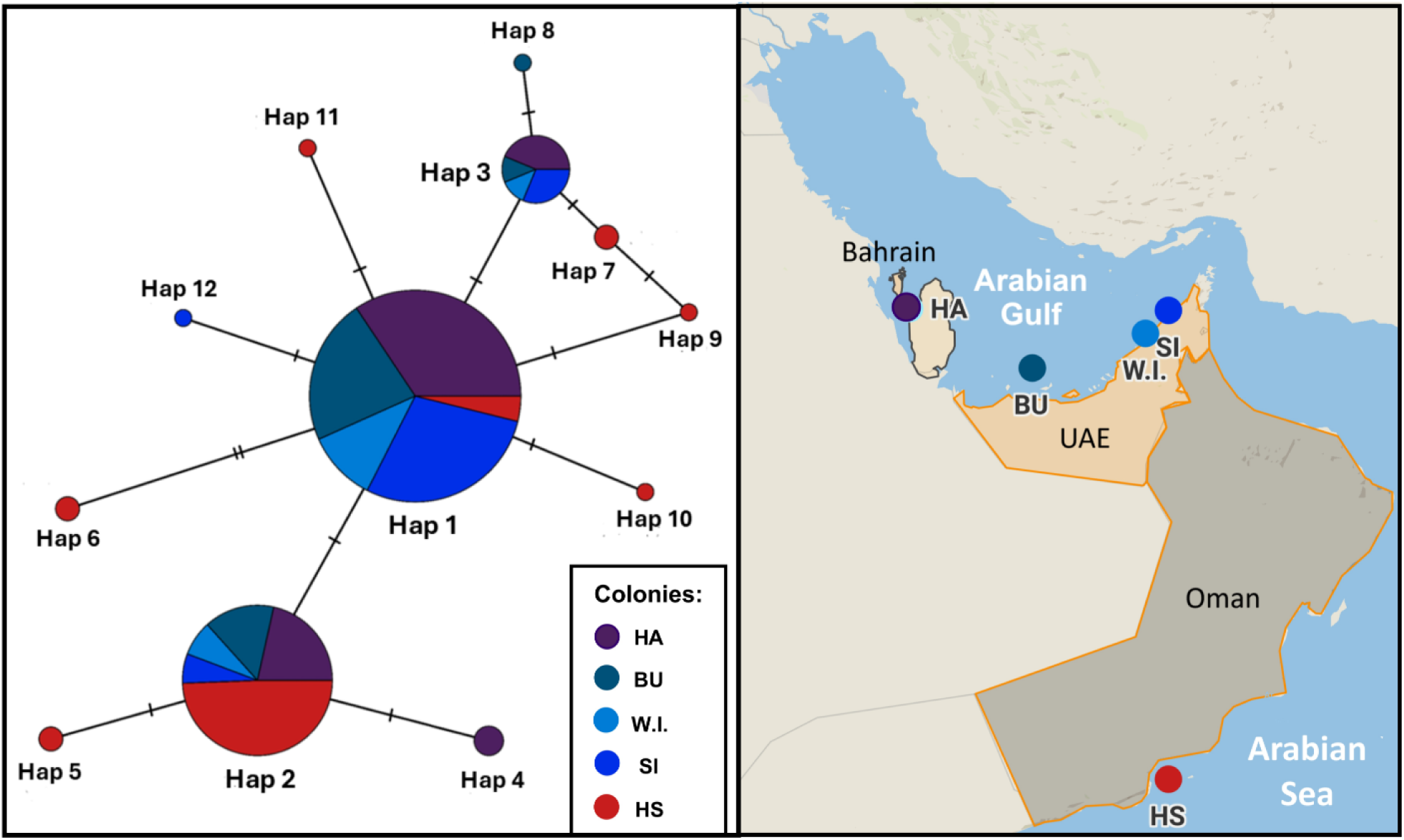
Median-joining network of 12 COI haplotypes across five Socotra cormorant colonies/sampling sites: HA = Hawar Island (Bahrain), BU = Butina Island (UAE), W.I. = World Islands (UAE), SI = Siniya Island (UAE), HS = Hasikiyah Island (Oman). Each pie-chart node represents one haplotype. Colors of nodes and pie-slices indicate the proportional number of individuals with a given haplotype per colony (site). Size of the nodes is proportional to frequency of each haplotype in sample. Black hatch-marks indicate number of mutations separating haplotypes. The network was constructed and visualized in PopART version 1.7 (Leigh and Bryant, 2015).

Structure clustered individuals into random mating groups, based on their haplotypes (COI) or genotypes (FIB7). For the COI dataset, both the Evanno method and Puechmaille estimators MedMedK and MedMeanK indicated *K* = 2 as the most likely number of clusters (see Figure S1 and S2A,C). The Structure bar plot for *K* = 2 showed individuals from the Arabian Gulf sites (HA, BU, W.I. and SI) predominantly assigned to one cluster, while individuals from the Arabian Sea site (HS) were mainly assigned to the second cluster (Figure 3A). The MaxMedK and MaxMeanK estimators supported *K* = 3 as the most likely number of COI clusters (Figure S2B,D). This pattern still assigned the Arabian Gulf colonies to a single cluster and HS (Arabian Sea) to a second cluster, with HS showing some admixture from a third cluster (Figure S3). For the FIB7, the Evanno method indicated *K* = 2 as the most likely number of clusters (Δ*K* = 184.64), and a log-likelihood values stabilizing from *K* = 2 (Figure S4). The Puechmaille method largely supported *K* = 2 (MedMedK, MaxMedK, MaxMeanK), although MedMeanK favored *K* = 1 (Figure S5), suggesting that the signal for *K* = 2 may reflect weak structure delineation. The Structure bar plot for *K* = 2 showed that individuals from all five colonies were predominantly assigned to a single cluster, with only minor admixture from the second cluster scattered among individuals—these clusters could not be assigned to any grouping of the colonies (Figure 3B). These results indicate that FIB7 exhibits very weak genetic differentiation among populations.

**FIGURE 3.**
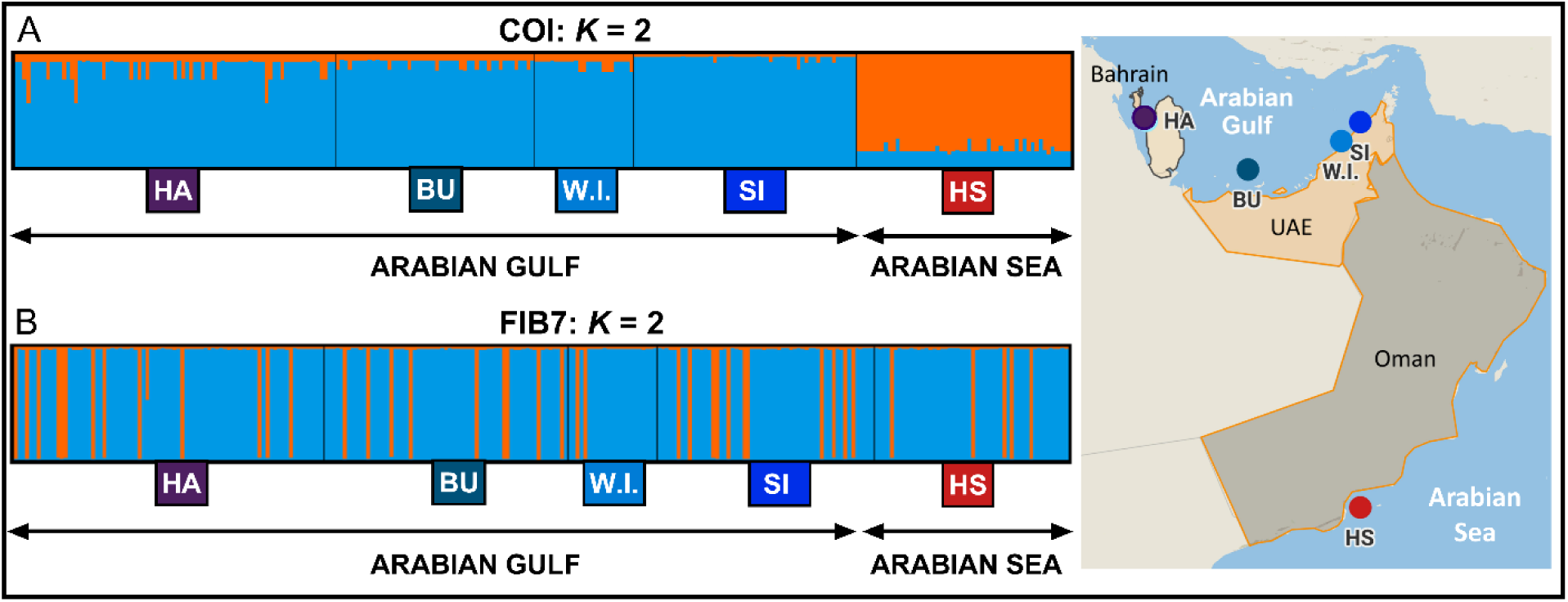
Individual cluster assignments for *K* = 2, estimated in Structure based on: (**A**) mitochondrial DNA cytochrome oxidase I (COI; 10 haploid SNPs) and (**B**) β-fibrinogen intron 7 (FIB7; 7 SNPs). Each vertical bar represents an individual, partitioned into colored segments corresponding to the estimated proportion of membership in each cluster. Sampling sites are indicated below: HA = Hawar Island (Bahrain), BU = Butina Island (UAE), W.I. = World Islands (UAE), SI = Siniya Island (UAE), HS = Hasikiyah Island (Oman). Cluster visualizations were generated using StructureSelector (Li and Liu 2018).

### Genetic diversity

The Arabian Gulf population harbored substantially lower mtDNA COI genetic diversity compared to the HS colony in the Arabian Sea (Table 2, Figure 4). A total of 6 private COI haplotypes were detected in the 54 individuals of the HS colony compared to only 3 private haplotypes for the entire Gulf population with 212 individuals; none of the Gulf colonies had more than one private haplotype (Table 2). The number of COI haplotypes detected in each colony is listed in Table S1 of the Supplementary Material. (Accession numbers of all COI haplotypes will be provided in Table S1 upon acceptance of manuscript.) Nucleotide diversity (π) for all colonies combined in the Arabian Gulf was only 0.00049 (SE 0.00001) compared to 0.00080 (SE 0.00001) for HS (Table 2, Figure 4). A two-tailed *t*-test comparing the subsamples of HS (6 x 9 sequences) and those of the Gulf population (23 x 9 sequencies) showed that the difference in COI π was highly significant (*t* = 3.306; *p* = 0.0027) (Table 2; Table S2). Also, comparisons of COI diversity between HS and the Gulf population, based on the subsampling approach, showed highly significant differences for the different number of haplotypes (*Na*: *t* = 4.03, *p* = 0.0004), Shannon’s information index (*I*: *t* = 3.35, *p* = 0.0024), nucleotide diversity (π: *t* = 3.31, *p* = 0.0027) and unbiased diversity (*uh*: *t* = 3.30, *p* = 0.0053) (Figure 4; Table 2; Table S2). For each of these diversity measures, the Gulf population displayed significantly lower diversity than HS (Table 2, Table S2). Likewise, statistical comparisons based on the mean values for *Na*, *I*, *uh* across the 10 SNPs, and derived from the full sample sets (HS: n = 54; Gulf: n = 212), showed significantly lower COI diversity for the combined Gulf population, despite featuring a much larger sample size (Figure 4; Table 2; Table S3). For all these diversity measures, the mean values for each of the four colonies in the Gulf were lower than that of HS in the Arabian Sea (Table 2). The only COI diversity measure that showed no significant difference between the Gulf and HS was haplotype diversity (*Hd*; Table 2; Tables S2, S3). We did not detect any significant difference for any of the diversity measures when comparing COI diversity between the four colonies inside the Arabian Gulf (HA, BU, W.I., SI) using non-parametric Kruskal-Wallis tests (Figure 4.; Table S4). The newly established colony on World Islands (W.I.) was the only one that did not have a private COI haplotype, but its genetic diversity values were somewhat intermediate compared to the other four colonies, including those in the Gulf (Table 2, Figure 4).

**FIGURE 4.**
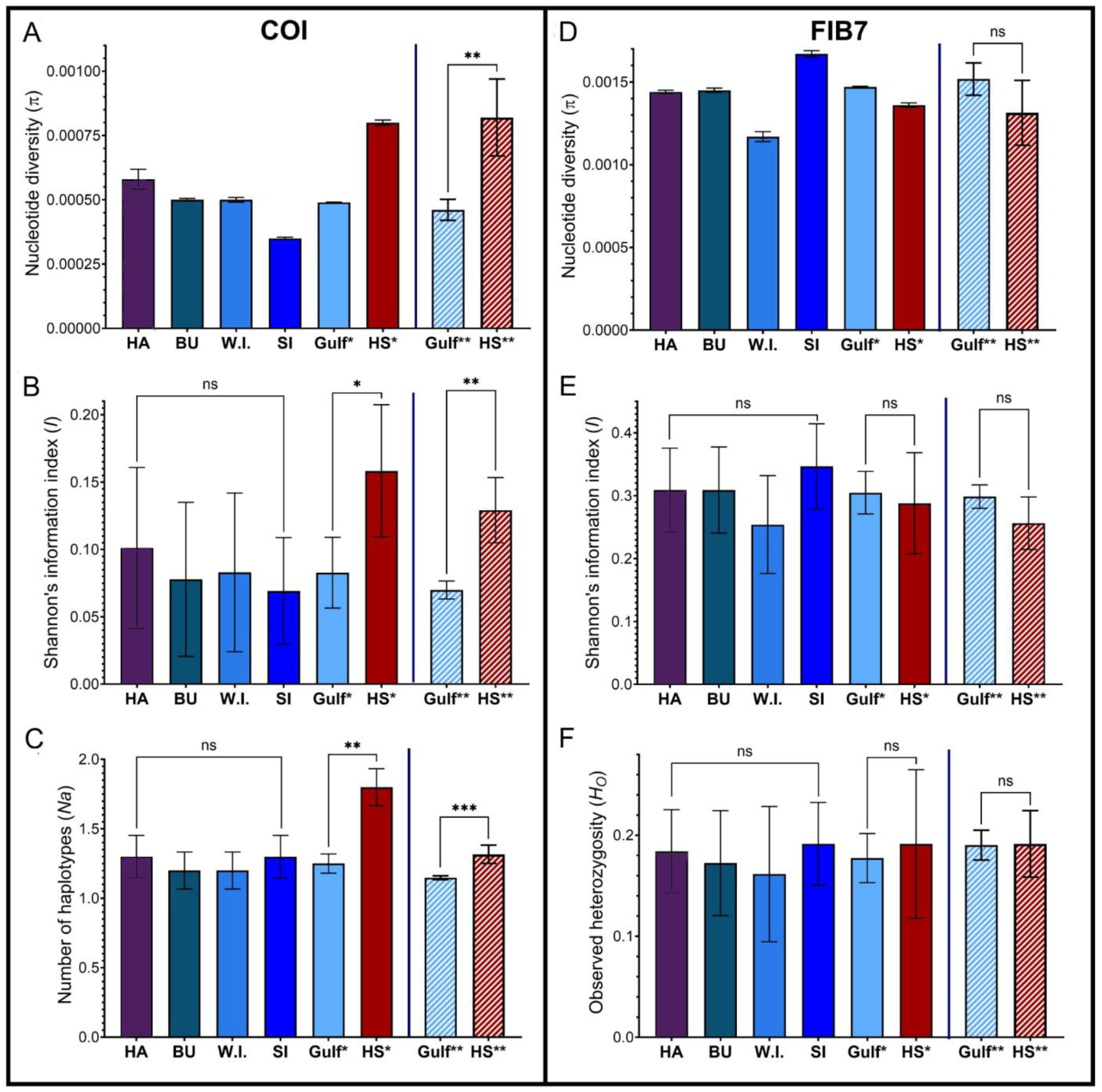
Genetic diversity measures for colonies (HA, BU, W.I., SI, HS) and regions (Gulf, HS). (**A-C**) measures based on mtDNA cytochrome oxidase I, (**D-F**) measures based on β-fibrinogen intron 7 (FIB7). Nesting colonies in Gulf: HA = Hawar, Bahrain; BU = Butina, UAE; W.I. = World Islands, UAE; SI = Siniya Island, UAE; nesting colony in Arabian Sea (HS = Hasikiyah Island, Oman). *Gulf & *HS: mean values and standard errors, based on estimates derived by Genalex for both regions across 10 and 7 SNPs for COI and FIB7, respectively. **Gulf & **HS: mean values and standard errors, based on estimates across 6 COI subsamples (each 9 sequences) for the Gulf colonies and 23 subsamples (each 9 sequences) for HS, and for β-fibrinogen intron 7 (FIB7) across 5 subsamples (each 10 sequences) for the Gulf and 22 subsamples (each 10 sequences) for HS. Whiskers show standard errors of the mean. sig. = statistical significance testing: ns = not significant (*p* ≥ 0.05), \**p* < 0.05; \*\**p* < 0.01.

**TABLE 2.**
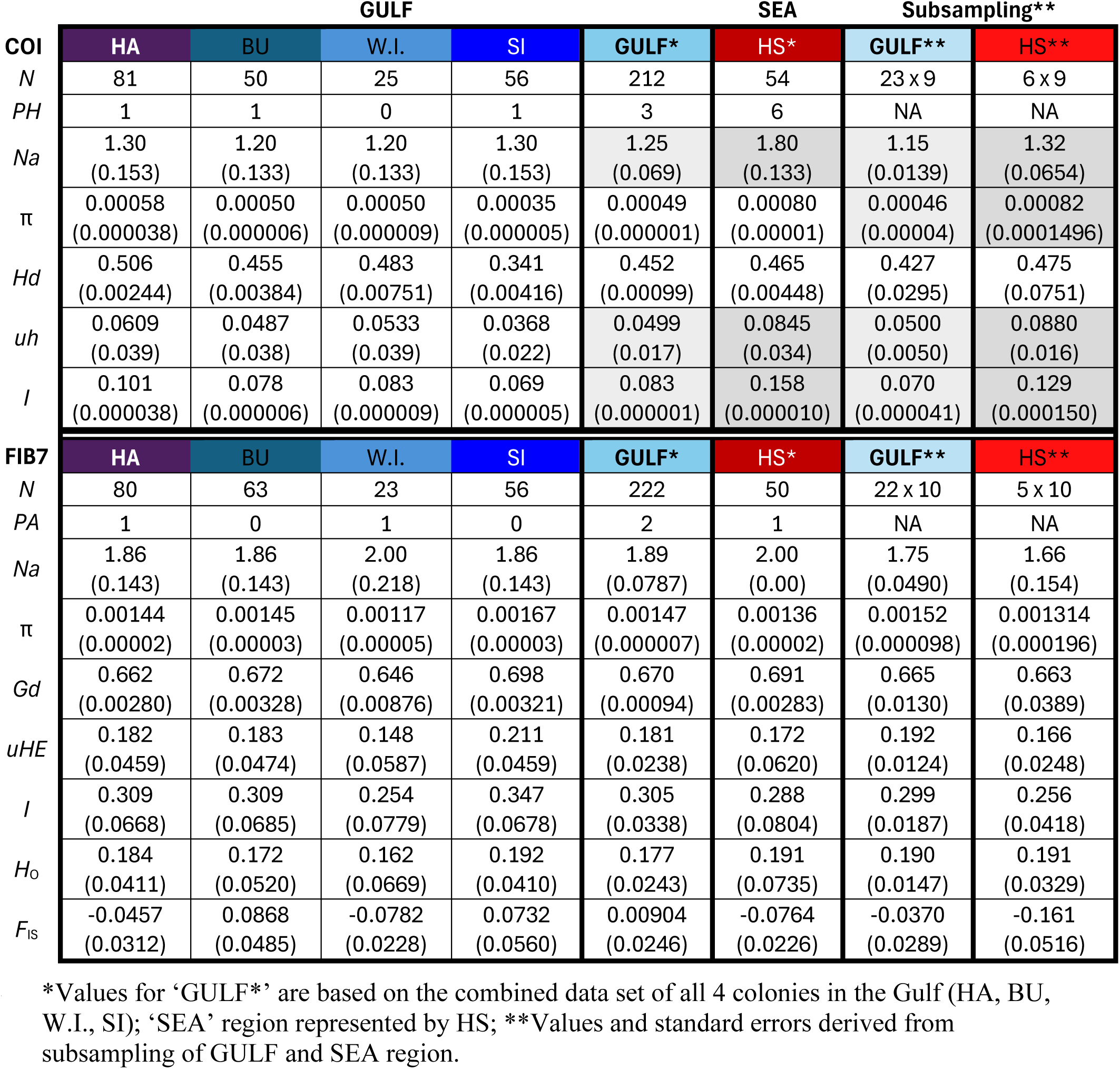
Genetic diversity measures, based on mtDNA cytochrome oxidase I (COI; top rows) and β-fibrinogen intron 7 (FIB7; bottom rows) for nesting colonies (HA = Hawar, Bahrain; BU = Butina, UAE; W.I. = World Islands, UAE; SI = Siniya Island, UAE; HS = Hasikiyah Island, Oman) and regions, ‘GULF’ & ‘SEA’ (represented by ‘HS’): *N* = sample size; *PH* = private haplotypes; *Na* = average number of different haplotypes (COI) or alleles (FIB7), π = nucleotide diversity; *Hd* = haplotype diversity; *uh* = unbiased diversity, *I* = Shannon’s information index; *Gd* = gene diversity; *uHe* = unbiased expected heterozygosity; *H*_O_ = observed heterozygosity; *F*_IS_ = fixation index. Standard errors of the mean are shown in parentheses. Shaded columns represent significant differences in values between regions (‘Gulf’ vs ‘HS’): values that are significantly greater represented by darker shades (Supplementary Material Tables S2, S3).

In contrast to the findings for COI, we did not detect any significant differences between the genetic diversity measure derived from FIB7 comparing the Gulf population with HS (Table 2; Figure 4; Tables S2, S3). Only 1 and 2 private FIB7 haplotypes (alleles) were detected in HS and the Gulf population, respectively (Table 2). The number of FIB7 haplotypes for each colony is listed in Table S1 of the Supplementary Material. (Accession numbers of all FIB7 haplotypes/alleles will be provided in Table S1.) Based on the subsampling approach, i.e. comparing subsamples with equal sample sizes, there were no significant differences between the Gulf and HS for any of the FIB7 diversity measures, namely π, *I*, gene diversity (*Gd*), unbiased expected heterozygosity (*uHe*), observed heterozygosity (*H*_O_), and the fixation index, *F*_IS_ (Table 2, Figure 4; Table S2). Likewise, no significant differences in diversity measures were detected when comparing the two populations (Gulf vs HS) for the full sample sets across the 7 FIB7 SNPs (Table 2; Figure 4, Table S3). The *F*_IS_ values, based on the entire sample size indicated a slight heterozygote excess (*F*_IS_ = − 0.0764; SE ± 0.0226) for HS, but not for the Gulf (*F*_IS_ = + 0.00904; SE ± 0.0246) (Table 2). Comparisons of diversity measures between the four colonies inside the Gulf (HA, BU, W.I., SI) showed no significant differences based on Kruskal-Wallis tests (Figure 4; Table S4). Of the four colonies inside the Arabian Gulf, the newly established W.I. colony had the lowest genetic diversity, based on nucleotide diversity (π), gene diversity (*Gd*), Shannon’s information index (*I*) and observed heterozygosity, but was the only colony in the Gulf with a private allele, and had the lowest sampling size (n = 23).

## DISCUSSION

### COI variation: well-connected population inside Gulf with some impediments to gene flow

This study presents the first characterization of population structure and genetic diversity of Socotra Cormorants in the region. The results, based on COI, suggest that the Socotra Cormorant colonies inside the Arabian Gulf (HA, BU, W.I., SI) form one large population that is isolated from Hasikiyah in the Arabian Sea (Table 1A,B, Figure 3A). The findings indicate high connectivity among Gulf colonies, with extensive gene flow inferred from COI, although a small but significant differentiation between HA and SI indicates subtle matrilineal structuring rather than complete panmixia. This is consistent with previous studies observing inter-colony movements, such as individuals from Siniya Island (SI) roosting with individuals from other colonies off the coast of Abu Dhabi, including Butina Island (BU), Um Qassar (sometimes called Umm Jassar Island), Umm Al Kurkum Islands, and islands off Dubai such as the World Islands site (W.I.) (Muzaffar et al. 2017b, Muzaffar 2020, Orben et al. 2025).

The only exception to this pattern of panmixia is a low but significant genetic differentiation found between cormorants of Hawar Island (HA), situated off Bahrain west of the Qatar Peninsula, and Siniya Island (SI) in the eastern part of the Gulf and (Figure 1). This may suggest subtle structuring within the Gulf, possibly driven by geographic distance or localized demographic processes. Given that Socotra Cormorants have been reported to fly long distances (e.g. Cook et al. 2016, Muzaffar et al. 2017a), and flyways are along the coast, it seems unlikely that the Qatar Peninsula itself represents a physical barrier to gene flow between those two colonies. However, gene flow could be indirectly impeded by water circulation pattens of the Arabian Gulf (Polikarpov et al. 2016): A frontal zone from the northern tip of Qatar effectively separates the waters between the Qatar Peninsula (Piontkovski et al. 2019), segregating plankton communities into distinct clusters along the frontal zones (Polikarpov et al. 2016). This segregation may partly explain the limited gene flow, inferred by COI variation, between SI and HA. Such local environment-dependent population structuring has been observed for other marine organisms, e.g. Common Bottlenose Dolphins (*Tursiops truncatus*) in the Black Sea, North Atlantic and Mediterranean Sea (Natoli et al. 2005, Moore et al. 2025), as well as seabirds (Friesen et al. 2007). However, the fact that significant COI *ΦST* and *Shua* values were only observed between Hawar (HA) and Siniiya (SI), the two most distant colonies sampled in the Gulf, and not between HA vs BU or HA vs W.I., suggests that this differentiation is more likely due to isolation by distance (IBD), rather than isolation by environment. In other words, the distance (ca. 480 km between HA and SI) may be a greater impediment to gene flow than environmental variation—otherwise we would also expect to observe genetic differentiation for HA vs BU or HA vs W.I. (Testing for IBD, e.g. using a Mantel test, is unlikely to address this due to the limited number of sampling sites and the fact that distances derived from coordinates, as used in a Mantel test, do not represent the actual flyway distances of the birds along the coastline.) Notably, the Structure bar plots, based on all Puechmaille estimators, as well as the Evanno method, assigned the Hawar (HA) colony to the same COI cluster as all the other Gulf colonies, including Siniya (SI) (Figure 3A; Supplementary Material Figure S3). Together, these findings suggest that the Socotra cormorant colonies within the Arabian Gulf are well-connected, representing one large population. It is possible, though, that the relatively low polymorphism of the genetic markers used, including COI, makes detection of fine-scale population structure of the Socotra cormorants in the Gulf impossible, and would require multiple polymorphic markers or a genomic approach.

### COI variation: Gulf population is isolated from Hasikiyah (Arabian Sea)

The strong genetic differentiation between Hasikiyah (HS) in the Arabian Sea the colonies within the Arabian Gulf, inferred by the variation of the COI locus (*ΦST* and *SHua* for each pairing: *p* < 0.001), suggests that Socotra Cormorants in the Gulf are genetically isolated from the Hasikiyah colony, and possibly other unsampled colonies outside the Gulf. The Structure analysis, based on COI, supports this pattern, with *K* = 2 clusters corresponding to the Gulf and Arabian Sea regions (Figure 3A), and an alternative *K* = 3 assignment (Figure S3), in which Gulf colonies remain a single cluster while HS shows some admixture. The significant differences in COI genetic diversity—the Gulf population exhibiting much lower diversity—underscore this isolation. Such isolation, may be a function of the relatively large distance (ca. 900 km between HS and SI, the two closest colonies across the Strait of Hormuz), and reflects the known distribution pattern of Socotra Cormorants in the region: there is no overlap between colonies inside the Arabian Gulf and those in the Arabian Sea (Jennings, 2010, BirdLife International 2019, Muzaffar et al. 2020). It is also consistent with the segregation of water masses and faunal communities on either side of the Strait of Hormuz (Piontkovski et al. 2019). The relatively large differentiation may also be explained by phylogeographic processes: The Arabian Gulf is a relatively recent Gulf that underwent flooding over a period of at least 18,000 years before present (b.p.) (Smith et al. 2022). The initial wave of flooding only covered a small portion of the northern extent of the Arabian Gulf. Further flooding until 12,000 years b.p. flooded the northwestern extent of the Arabian Gulf. It was not until less than 8000 years b.p., that the Gulf achieved its current extent, with all of the southern extent comprising the United Arab Emirates shoreline being inundated (Smith et al. 2022). It is conceivable that small populations of Socotra Cormorants expanded into the Arabian Gulf early in its history and remained isolated from those in the Arabian Sea. These small populations, with relatively lower genetic diversity, presumably were founders of the early Arabian Gulf population that spread and proliferated over the 18,000-year period within the Gulf. The limited gene flow and relatively low COI diversity of Socotra Cormorants inside the Gulf is consistent with this hypothesis. On the other hand, the parent population in the Arabian Sea may have been more diverse and presumable expanded south towards the Gulf of Aden and the Red Sea. Although the COI-based signal of differentiation between the Gulf Socotra cormorants and the HS colony appears strong, further investigation using additional genetic markers would be beneficial. The inclusion of additional markers would help elucidate fine-scale population structure and strengthen the confidence of inferences regarding population structure, gene flow, and phylogeographic processes.

### Gulf Socotra Cormorants harbor relatively low genetic diversity

Our study also showed relatively low genetic diversity for this threatened seabird species across both genetic markers, mtDNA COI and FIB7 (Table 2). Particularly, COI diversity was comparatively low for the Gulf population. For instance, all colonies inside the Gulf combined harbored only 3 private COI haplotypes compared to 6 private haplotypes in the Hasikiyah colony alone. Also, both the subsampling approach and the statistical analyses comparing diversity measures across the SNPs showed that the Gulf population has significantly lower COI genetic diversity compared to Hasikiyah in the Arabian Sea (Table 2, Figure 4; Supplementary Material Table S2, Table S3). This relatively low COI diversity may be atypical for a population that shows high connectivity among most Gulf colonies (based on both nuclear and mitochondrial markers). Large, well-connected populations typically harbor greater genetic diversity compared to smaller, fragmented ones (Frankham 2010, Grueber and Jamieson 2011, Manlik et al. 2019a). Despite the fact that the Socotra cormorants inside the Gulf represent the most abundant aggregation of this species, the low diversity in the Gulf may reflect historical isolation and limited gene flow beyond the Gulf, which are key drivers of genetic erosion (Frankham 2010, Méndez et al. 2014). Moreover, low genetic diversity is associated with reduced fitness and a diminished capacity to adapt to environmental changes (Reed and Frankham 2003, Spielman et al. 2004, Frankham 2010). Therefore, our findings that the Socotra Cormorants inside the Arabian Gulf, display relatively low genetic diversity and restricted gene flow with other colonies outside the Gulf, raise concerns for the conservation of this Gulf population.

The relatively new colony on the man-made World Islands site (W.I.) displayed the lowest FIB7-diversity among the five colonies for almost all of the diversity measures (Table 2, Figure 4). This comparatively low FIB7 diversity may reflect a founder’s effect, which typically leads to a reduction in genetic variation when only a small number of individuals establish a new population (Templeton 1980, Star and Spencer 2013). In particular π, which is relatively insensitive to sample size compared to some of the other measures was substantially lower for W.I. compared to all other colonies. Hence, it is feasible that this relatively low FIB7-diversity of W.I. may reflect genetic drift via the founder’s effect. The W.I. colony was also the only colony without a private COI haplotype. However, COI diversity measures were somewhat intermediate for W.I. compared to the other colonies. A founder’s effect assumes that only a small number of individuals initiate a new population, which may or may not be the case for W.I. The relatively low sample size for W.I. makes subsampling unfeasible for additional statistical analyses to allow for a better comparison.

### Marker-specific differences

Our findings with respect to genetic differentiation and genetic diversity based on the two genetic markers are notably different. While no population structure was detected for the nuclear marker FIB7, the results for mitochondrial DNA COI revealed significant differentiation between the Socotra Cormorants in the Arabian Gulf and those from the Hasikiyah colony in the Arabian Sea. Likewise, FIB7 did not reveal any significant differences in genetic diversity between the two regions, whereas COI diversity was significantly lower for the Gulf population. The observation that genetic differentiation was inferred by COI and not FIB7 could be reflecting sex-specific dispersal. Given that mitochondrial DNA is maternally inherited, any measure of differentiation based on mtDNA only reflects dispersal of females, while variation in the biparentally inherited nuclear DNA reflects dispersal of both sexes. Hence, the significant differentiation based on COI (but not FIB7) might indicate that (compared to males) females show greater site fidelity, adhering to colonies on either side of the Strait of Hormuz. Such sex-specific dispersal is not commonly observed in seabirds, but is consistent with a study on Great Cormorants (*Phalacrocorax carbo*), which indicated that females showed greater natal site fidelity than males (Schjørring 2001). Males in that study were shown to disperse farther from natal breeding sites compared to females. However, there is no evidence to indicate that dispersal of Socotra Cormorants is sex-specific—males and females have been reported to disperse similar distances and to similar regions post-breeding (Muzaffar et al. 2017a). An alternative explanation for the discrepancy in results between the two markers may be that the very low polymorphism of the non-coding FIB7 pre-empts the inference of population structure. Notably, a meta-analysis showed no correlation between mtDNA and nuclear microsatellite diversity across 54 bird species (Schmidt et al. 2023), so the discrepancy between the results we obtained for mtDNA versus FIB7 may not be too surprising. It would be desirable to reassess population structure, based on additional, more polymorphic loci, e.g., microsatellite loci, or ideally use a genomic approach to infer population structure of the Socotra Cormorants in their distribution range. In particular, Structure analysis is typically conducted on datasets with multiple, highly polymorphic microsatellite loci (e.g. Manlik et al. 2019a, Graham et al. 2022, Hernández-Soto et al. 2023), which provide greater resolution for detecting fine-scale population structure. Therefore, to achieve a more robust understanding of population structure of the Socotra Cormorants, further investigation using multiple genetic loci or genomic data (e.g. ddRAD sequencing) should be conducted.

## Conclusion

This study provides the first characterization of population structure and genetic diversity of the regionally endemic Socotra Cormorants. Our findings suggest that the Socotra Cormorants inside the Arabian Gulf form a large population that appears to be isolated from the Socotra Cormorants on the Hasikiyah Islands breeding site (HA) in the Arabian Sea. Colonies within the Gulf are well connected, exhibiting extensive gene flow, with the exception of the two most distant sites in the Gulf, Hawar (HA) and Siniya Island (SI), which display more limited gene flow, based on COI variation. This isolation of the Gulf Socotra Cormorants from Hasikiyah in the Arabian Sea is consistent with the known distribution and flight patterns of Socotra Cormorants and the phylogeography of the region. We thus support previous calls (BirdLife International 2019, Muzaffar et al. 2017a) for treating the Gulf and Arabian Sea Socotra Cormorants as separate evolutionarily significant units (ESUs) (Moritz 2004). The comparatively low mtDNA genetic diversity for the Socotra Cormorant population inside the Gulf might be due to its relative isolation from colonies outside the Gulf. The low genetic diversity in combination with the relative isolation of the Gulf Socotra Cormorants also raises concerns regarding their long-term viability, as these factors may be associated with reduced fitness and a diminished evolutionary potential to adapt to environmental change. Future research should focus on genomic analyses to refine population structure and assess adaptive potential, as well as conservation strategies to mitigate risks associated with genetic erosion.

## Supplementary material

Additional supporting information may be found online in the Supplementary Material:

**FIGURE S1**: Optimum *K* for COI dataset (Evanno method): Mean posterior probabilities (Ln P(D)) and Δ*K* values (Evanno et al. 2005) calculated for COI (Structure analysis)

**FIGURE S2**: Optimum *K* for COI dataset, based on Puechmaille method (STRUCTURESELECTOR): Median-based (MedMedK and MaxMedK) and mean-based (MedMeanK and MaxMeanK) estimators for the optimal number of clusters (*K*).

**FIGURE S3**: Alternative cluster assignment for COI, *K* = 2, estimated in STRUCTURE

**FIGURE S4**: Optimum *K* for FIB7 dataset (Evanno): Mean posterior probabilities (Ln P(D)) and Δ*K* values (Evanno et al. 2005) calculated for FIB7 (Structure analysis)

**FIGURE S5**: Optimum *K* for FIB7 dataset, based on Puechmaille method (STRUCTURESELECTOR): Median-based (MedMedK and MaxMedK) and mean-based (MedMeanK and MaxMeanK) estimators for the optimal number of clusters (*K*).

**Table S1**: List of COI & FIB7 haplotypes/alleles, relative frequencies per colony, and associated accession numbers

**Table S2**: Two-tailed *t*-tests and Mann-Whitney *U* test comparing genetic diversity values between subsamples of the Arabian Gulf and Hasikiyah of the Arabian Sea.

**Table S3**: Two-tailed Mann-Whitney *U* tests comparing genetic diversity values between Arabian Gulf and Hasikiyah of the Arabian Sea, derived from entire sample sets.

**Table S4**: Kruskal-Wallis statistics comparing genetic diversity values between the four colonies inside the Gulf

## Acknowledgments

We thank the United Arab Emirates University (UAEU) for the generous funding that supported this study, including a Start-Up grant and a Sustainability Development Goals (SDG) grant to Oliver Manlik (see ‘Funding’ below). Additionally, open access publishing was facilitated by UAEU. We are thankful for the Environment Agency—Abu Dhabi (EAD) for providing us with essential logistic assistance and permitting to access and collect samples from Butina Island as part of the Marawah Marine Biosphere Reserve, the Umm Al Quwain Municipality for the permission to collect sample from Siniya Island, and the Dubai Environment and Climate Change Authority for permitting sample collection from the World Islands. We also thank the Arab Regional Center for World Heritage for administrative support and assisting in issuing all relevant permits, the Supreme Council for Environment, Bahrain, for issuing the environmental permits and logistic support, e.g. proving a boat & research assistants, as well as the Southern Tourism Company for logistics associated with accommodation and transport in Bahrain. Also, thanks to Reneco International Wildlife Consultants LTD, which supported Noura Almansoori during the time of writing her thesis and analyses that led to this paper.

## Ethics statement

All protocols for this study were approved by the United Arab Emirates University (UAEU) Animal Ethics committee with the protocol number ERA_2019_5969. Blood samples from Socotra Cormorants were collected under the Environment Agency – Abu Dhabi (EAD) permit #13676. The Arab Regional Center for World Heritage, Bahrain and the Supreme Council for Environment, Bahrain issued relevant environmental survey permits, as well as export permits. No animal was sacrificed or removed from its natural habitat for the purpose of this study.

## Author contributions

NMA: Conceptualization; methodology (lead); software; investigation; validation; formal analysis; visualization; resources; writing original draft; writing - review & editing. HR: Conceptualization; methodology; data curation; investigation; project administration; resources; writing - review & editing. SBM: Conceptualization; methodology; data curation; investigation; supervision; project administration; resources; writing - review & editing. DBHC: methodology; software; validation; formal analysis; visualization; writing - review & editing. AN: methodology; data curation; investigation; validation; writing - review & editing. MA: methodology; data curation; resources; writing - review & editing. HN: data curation; resources; writing - review & editing. AK: data curation; resources; writing - review & editing. FAH: methodology; investigation. LSRA: methodology; investigation. MMBA: methodology; investigation. FMAD: methodology; data curation; writing - review & editing. OM: Conceptualization (lead); methodology; software; data curation; investigation; validation; formal analysis; supervision (lead); funding acquisition, writing - review & editing.

## Data availability

Additional data are available in the Supplementary Material—see above. Individual DNA sequences (FIB7 and COI), as well as sequence alignments, will be uploaded on NCBI GenBank, and corresponding accession numbers for the new Socotra Cormorant haplotypes and alleles of this study will be provided upon acceptance of manuscript (‘TBA’).

