## Supplementary Material for "Socotra Cormorants in the Arabian Gulf represent a large, but isolated population with low genetic diversity"

**SUPPLEMENTARY FIGURES:**

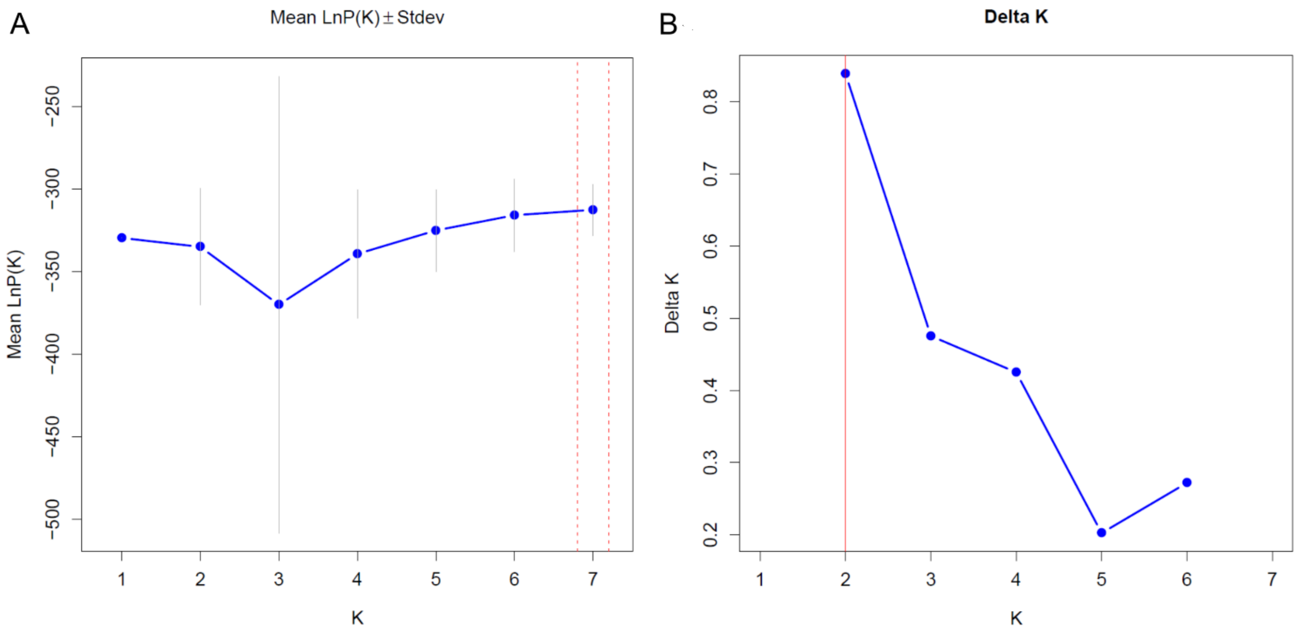


**FIGURE S1.** (**A**) Mean posterior probabilities (Ln P(D)) and (**B**) Δ*K* values (Evanno et al. 2005) calculated across 25 replicate runs for *K* = 1–7 using the Bayesian clustering method implemented in Structure for the COI dataset and summarized using StructureSelector. The red line indicates the best *K* value suggested by the Evanno method (Evanno et al. 2005). Mean Ln P(D) values increased with *K* and peaked at *K* = 7, as indicated by the dashed vertical lines (panel **A**), whereas the Evanno method identified *K* = 2 as the most likely number of clusters based on Δ*K* (panel **B**). This discrepancy is common because Ln P(D) tends to rise with *K*, whereas Δ*K* detects the strongest hierarchical level of structure.

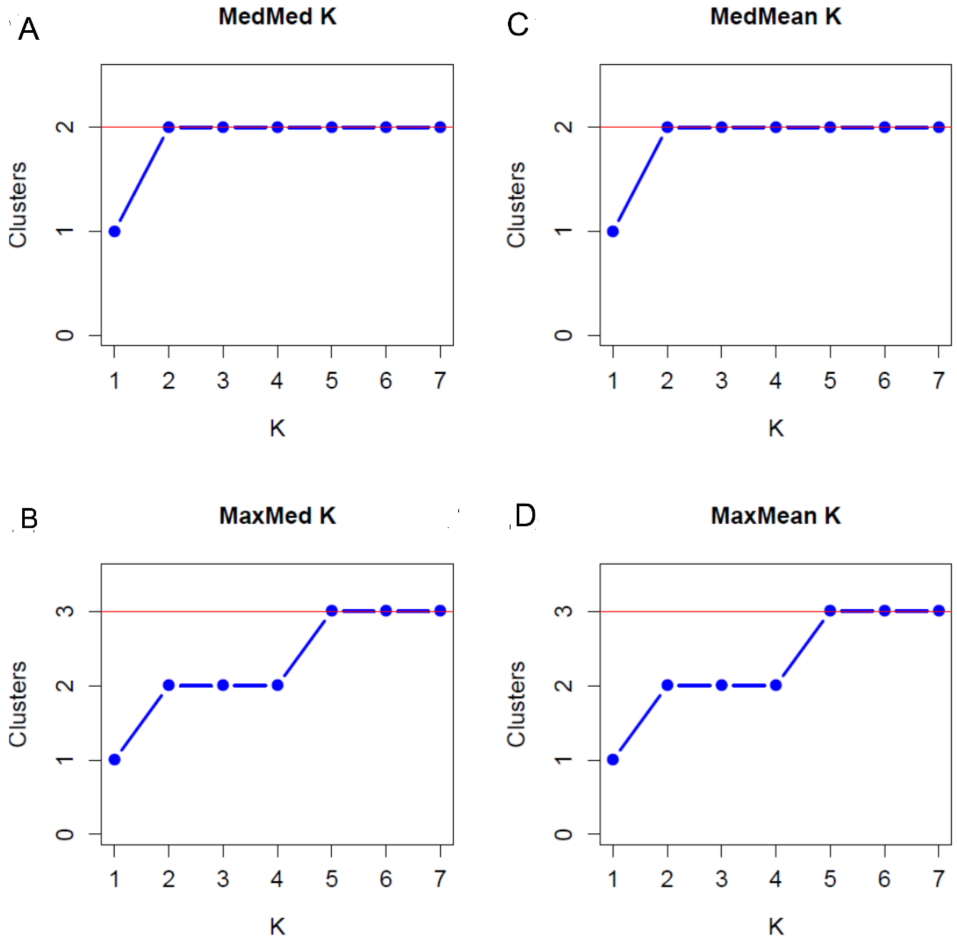


**FIGURE S2**. (**A**, **B**) Median-based (MedMedK and MaxMedK) and (**C**, **D**) mean-based (MedMeanK and MaxMeanK) estimators for the optimal number of clusters (*K*) (Evanno et al. 2005) for the COI dataset, calculated using STRUCTURESELECTOR (Li and Liu 2018). The red line indicates the best number of clusters identified by each estimator.

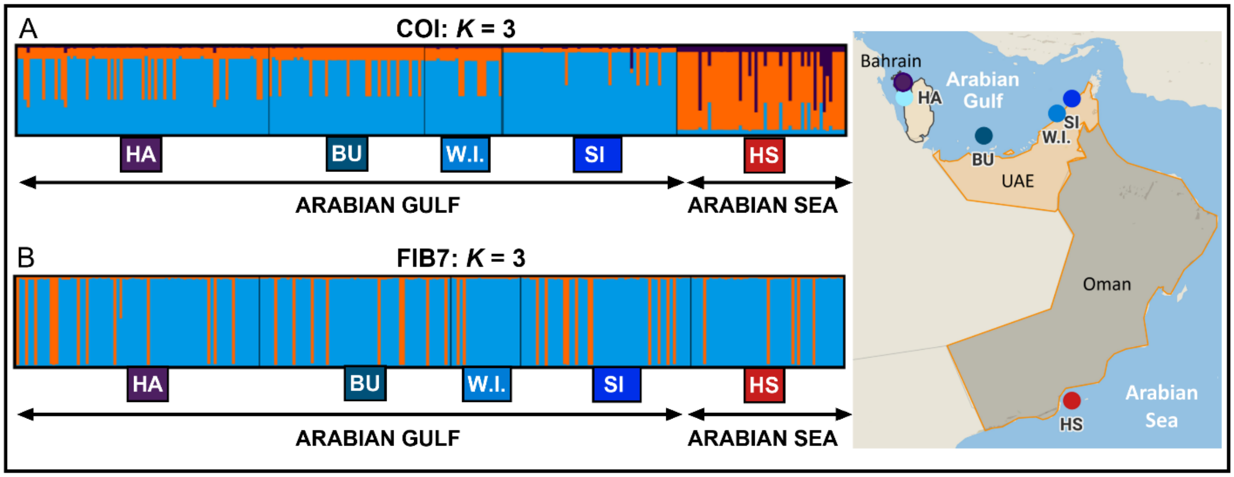


**FIGURE S3**. Alternative COI cluster assignments for $K = 3$, estimated in STRUCTURE based on: (**A**) mitochondrial DNA cytochrome oxidase I (COI; 10 haploid SNPs) and (**B**) β-fibrinogen intron 7 (FIB7; 7 SNPs). (Structure assignment for COI $K = 2$ shown in Fig. 3 of main text.) Each vertical bar represents an individual, partitioned into colored segments corresponding to the estimated proportion of membership in each cluster. Sampling sites are indicated below. Sampling sites are indicated below: HA = Hawar Island (Bahrain), BU = Butina Island (UAE), W.I. = World Islands (UAE), SI = Siniya Island (UAE), HS = Hasikiyah Island (Oman). Cluster visualizations were generated using STRUCTURESELECTOR (Li and Liu 2018).

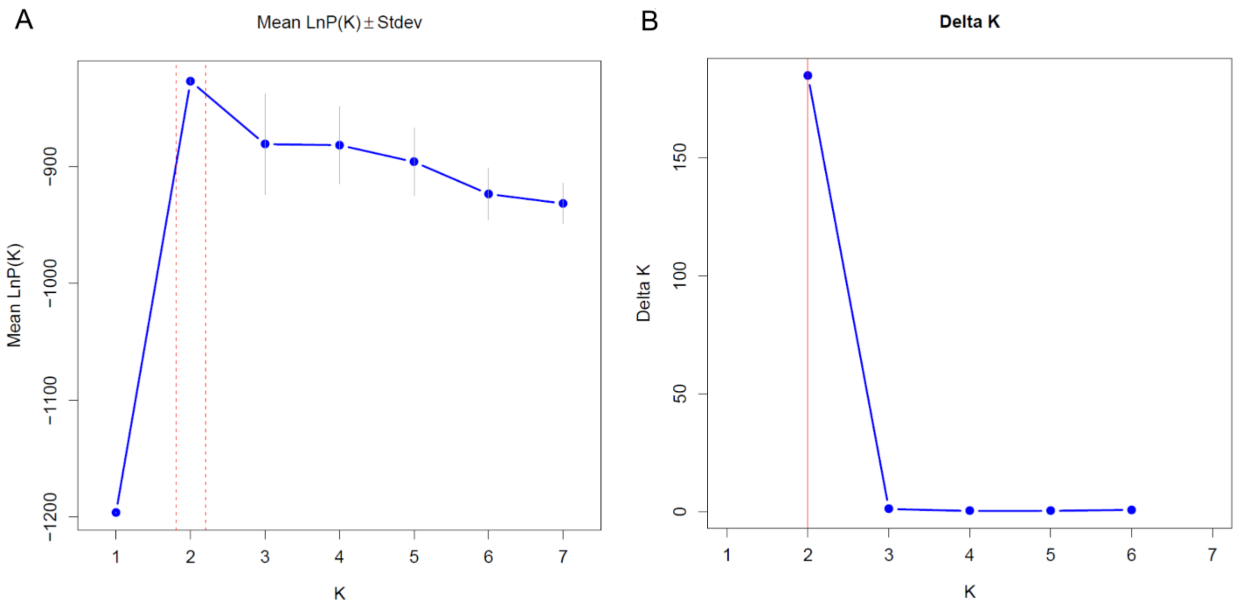


**Figure S4. (A)** Mean posterior probabilities (Ln P(D)) and **(B)** $\Delta K$ values (Evanno et al. 2005) calculated across 25 replicate runs for $K = 1–7$ using the Bayesian clustering method implemented in STRUCTURE for the FIB7 dataset and summarized using STRUCTURESELECTOR. . The dashed vertical line in panel **A** indicates the *K* value with the highest mean Ln P(D), whereas the red line in panel **B** indicates the best *K* value suggested by the Evanno method (Evanno et al. 2005).

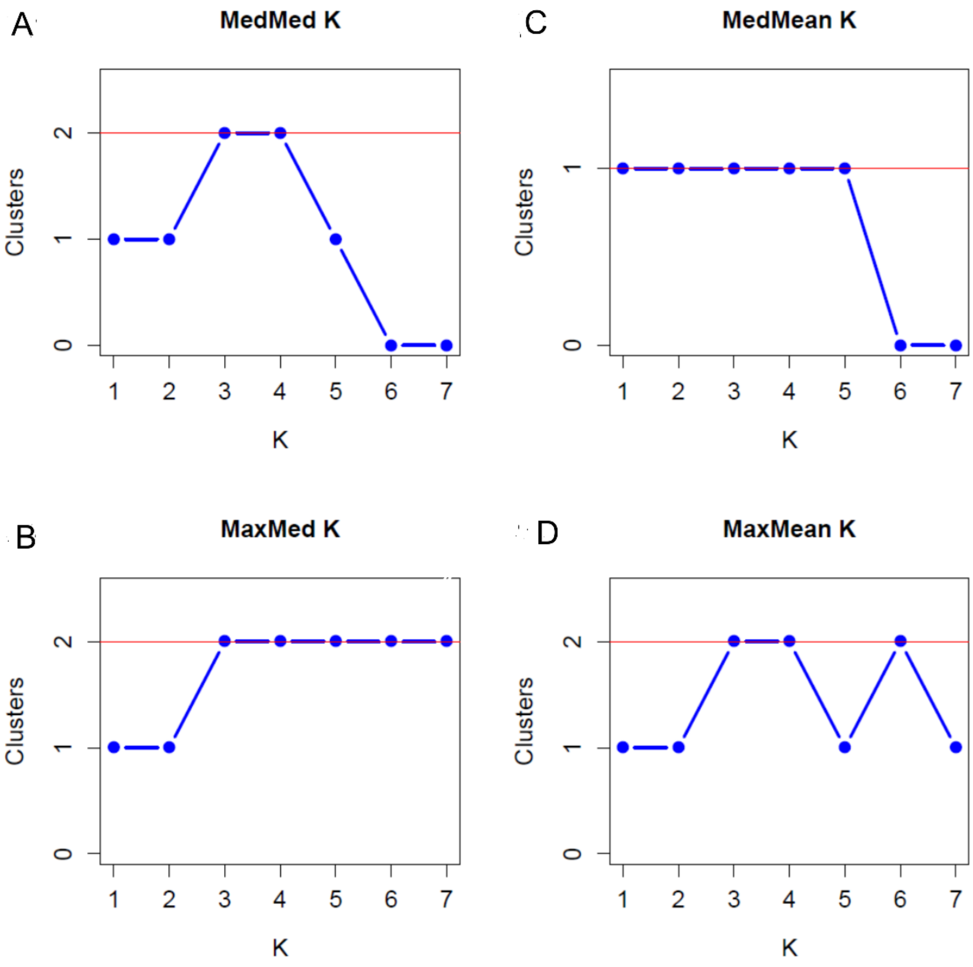


**Figure S5. (A, B)** Median-based (MedMedK and MaxMedK) and **(C, D)** mean-based (MedMeanK and MaxMeanK) estimators for the optimal number of clusters (*K*) (Evanno et al. 2005) for the FIB7 dataset, calculated using StructureSelector (Li and Liu 2018). The red line indicates the best number of clusters identified by each estimator.

**SUPPLEMENTARY TABLES:**

**TABLE S1**: List of COI & FIB7 haplotypes/alleles, relative frequencies per colony, and associated accession numbers

**TABLE S2**: Two-tailed *t*-tests and Mann-Whitney *U* test comparing genetic diversity values between subsamples of the Arabian Gulf and Hasikiyah of the Arabian Sea.

**TABLE S3**: Two-tailed Mann-Whitney *U* tests comparing genetic diversity values between Arabian Gulf and Hasikiyah of the Arabian Sea, derived from entire sample sets.

**TABLE S4**: Kruskal-Wallis statistics comparing genetic diversity values between the four colonies inside the Gulf

**TABLE S1.** Haplotypes for COI and FIB7 (alleles) of Socotra Cormorants (*Phalacrocorax nigogularis*) in Arabian Gulf ('Gulf') and Hasikiyah ('HS'), inferred by DNASP (Librado and Rozas, 2009). Haplotypes (alleles) are tabulated in order of frequency across all colonies (Hap 1 to Hap 12 for COI, top rows; Hap 1-7 for FIB7; lower rows). Occurrence of haplotypes/alleles in either Gulf, HS or both is indicated in the 'Location' column. Also listed are the closest BLAST matches. Note: Hap 2 is identical to *P. nigrogularis* sequences available as accession number KM066529 (Kennedy and Spencer 2014) in GenBank. Accession numbers for the new Socotra Cormorant haplotypes and alleles of this study will be provided upon acceptance of manuscript ('TBA'). Alignments of COI haplotypes and FIB7 alleles will be uploaded on GenBank with accession numbers XXXX and XXXX, respectively. The number of haplotypes (COI) and alleles (FIB7) for each colony (HA = Hawar, Bahrain; BU = Butina, UAE; W.I. = World Islands, UAE; Siniya Island, UAE; HS = Hasikiyah Island, Oman) are listed in the five columns on the five right.

| COI | Accession # | Species | Study | BLAST closest species | HA | BU | W.I. | SI | HS |
|---|---|---|---|---|---|---|---|---|---|
| **Hap 1** | TBA | *P. nigrogularis* | This study | *P. nigrogularis* (KM066529) | 54 | 35 | 17 | 45 | 6 |
| **Hap 2** | KM066529** | *P. nigrogularis* | Kennedy and Spencer 2014 | *P. nigrogularis* (100% match: KM066529) | 17 | 12 | 6 | 5 | 39 |
| **Hap 3** | TBA | *P. nigrogularis* | This study | *P. nigrogularis* (KM066529) | 7 | 2 | 2 | 5 | 0 |
| **Hap 4** | TBA | *P. nigrogularis* | This study | *P. nigrogularis* (KM066529) | 3 | 0 | 0 | 0 | 0 |
| **Hap 5** | TBA | *P. nigrogularis* | This study | *P. nigrogularis* (KM066529) | 0 | 0 | 0 | 0 | 2 |
| **Hap 6** | TBA | *P. nigrogularis* | This study | *P. nigrogularis* (KM066529) | 0 | 0 | 0 | 0 | 2 |
| **Hap 7** | TBA | *P. nigrogularis* | This study | *P. nigrogularis* (KM066529) | 0 | 0 | 0 | 0 | 2 |
| **Hap 8** | TBA | *P. nigrogularis* | This study | *P. nigrogularis* (KM066529) | 0 | 1 | 0 | 0 | 0 |
| **Hap 9** | TBA | *P. nigrogularis* | This study | *P. nigrogularis* (KM066529) | 0 | 0 | 0 | 0 | 1 |
| **Hap 10** | TBA | *P. nigrogularis* | This study | *P. nigrogularis* (KM066529) | 0 | 0 | 0 | 0 | 1 |
| **Hap 11** | TBA | *P. nigrogularis* | This study | *P. nigrogularis* (KM066529) | 0 | 0 | 0 | 0 | 1 |
| **Hap 12** | TBA | *P. nigrogularis* | This study | *P. nigrogularis* (KM066529) | 0 | 0 | 0 | 1 | 0 |

| FIB7 | Accession # | Species | Study | BLAST closest species | HA | BU | W.I. | SI | HS |
|---|---|---|---|---|---|---|---|---|---|
| **Hap 1** | TBA | *P. nigrogularis* | This study | *P. neglectus* (KM066288) | 79 | 60 | 22 | 49 | 37 |
| **Hap 2** | TBA | *P. nigrogularis* | This study | *P. neglectus* (KM066288) | 41 | 31 | 16 | 29 | 37 |
| **Hap 3** | TBA | *P. nigrogularis* | This study | *P. neglectus* (KM066288) | 26 | 25 | 6 | 22 | 20 |
| **Hap 4** | TBA | *P. nigrogularis* | This study | *P. neglectus* (KM066288) | 13 | 10 | 1 | 12 | 5 |
| **Hap 5** | TBA | *P. nigrogularis* | This study | *P. neglectus* (KM066288) | 0 | 0 | 0 | 0 | 1 |
| **Hap 6** | TBA | *P. nigrogularis* | This study | *P. neglectus* (KM066288) | 1 | 0 | 0 | 0 | 0 |
| **Hap 7** | TBA | *P. nigrogularis* | This study | *P. neglectus* (KM066288) | 0 | 0 | 1 | 0 | 0 |

**TABLE S2.** Two-tailed *t*-tests and Mann-Whitney *U* test* comparing genetic diversity values between subsamples of the Arabian Gulf ('GULF') and Hasikiyah ('HS') of the Arabian Sea. Comparisons are based on mean values for mtDNA cytochrome oxidase I (COI; top rows) across 23 subsamples (each 9 sequences) for the GULF colonies and 6 subsamples (each 9 sequences) for HS, and for β-fibrinogen intron 7 (FIB7; bottom rows) across 22 subsamples (each 10 sequences) for the GULF colonies and 5 subsamples (each 10 sequences) for HS. $\pi$ = nucleotide diversity, $I$ = Shannon's information index; *uh* = unbiased diversity (COI), *Na* = number of different haplotypes (COI) or alleles (FIB7), *Hd* = haplotype diversity (COI), *Gd* = gene diversity (FIB7), *uHe* = unbiased expected heterozygosity (FIB7), $H_O$ = observed heterozygosity (FIB7), $F_{IS}$ = fixation index (FIB7). sig. = statistical significance testing for *t*-tests and Mann-Whitney *U* test: ns = not significant ($p \geq 0.05$), *$p < 0.05$; **$p < 0.01$; ***$p < 0.001$. Mean values and standard errors of the means (in parentheses) across subsamples are listed for each diversity measure for GULF and HS—these are also tabulated in Table 2 of the main text.

| **COI** | *t* | *p* | sig. | **GULF** | HS |
|---|---|---|---|---|---|
| $\pi$ | 3.306 | 0.0027 | ** | 0.00046 (0.00004) | 0.00082 (0.0001496) |
| $I$ | 3.348 | 0.0024 | ** | 0.070 (0.000041) | 0.129 (0.000150) |
| *uh* | 3.303 | 0.0053 | ** | 0.050 (0.0050) | 0.088 (0.016) |
| *Na* | 4.028 | 0.0004 | *** | 1.15 (0.0139) | 1.32 (0.0654) |
| *Hd* | 0.6914 | 0.4952 | ns | 0.427 (0.0295) | 0.475 (0.0751) |

| **FIB7** | *t* | *p* | sig. | **GULF** | HS |
|---|---|---|---|---|---|
| π | 0.9017 | 0.3758 | ns | 0.00152 (0.000098) | 0.001314 (0.000196) |
| $I$ | 0.9638 | 0.3444 | ns | 0.299 (0.0187) | 0.256 (0.0418) |
| *uHe* | 0.9060 | 0.3736 | ns | 0.192 (0.0124) | 0.166 (0.0248) |
| *Gd* | 0.04268 | 0.9663 | ns | 0.665 (0.0130) | 0.663 (0.0389) |
| $H_O$ | 0.03394 | 0.9732 | ns | 0.190 (0.0147) | 0.191 (0.0329) |
| $F_{IS}$ | 1.8820 | 0.0715 | ns | -0.0370 (0.0289) | -0.161 (0.0516) |

| **FIB7** | *U** | *p* | sig. | **GULF** | HS |
|---|---|---|---|---|---|
| *Na** | 51.5 | 0.7117 | ns | 1.75 (0.0490) | 1.66 (0.154) |

*Distribution for FIB7 number of alleles (*Na*) significantly departed from normality and could not be transformed to a normal distribution, so statistical comparison is based on non-parametric Mann-Whitney *U* test.

**TABLE S3.** Two-tailed Mann-Whitney *U* tests comparing genetic diversity values between the Arabian Gulf ('GULF') and Hasikiyah ('HS') of the Arabian Sea, derived from entire sample sets. Comparisons are based on mean values estimated with GenAlEx for cytochrome oxidase I (COI; top rows) and β-fibrinogen intron 7 (FIB7; bottom rows) across 10 and 7 SNPs, respectively. *I* = Shannon's information index, *Na* = number of different haplotypes (COI) or alleles (FIB7), *uh* = unbiased diversity (COI), *uHe* = unbiased expected heterozygosity (FIB7), $H_O$ = observed heterozygosity (FIB7), $F_{IS}$ = fixation index (FIB7). sig. = statistical significance testing Mann-Whitney *U* test: ns = not significant ($p \geq 0.05$), $*p < 0.05$; $**p < 0.01$. Mean values and standard errors of the means (in parentheses) across the SNPs are listed for each diversity measure for GULF and HS—these are also tabulated in Table 2 of the main text.

| **COI** | *U* | *p* | sig. | **GULF** | HS |
|---|---|---|---|---|---|
| *I* | 111 | 0.0143 | * | 0.083 (0.000001) | 0.158 (0.000010) |
| *Na* | 90 | 0.0023 | ** | 1.25 (0.069) | 1.80 (0.133) |
| *uh* | 110.5 | 0.0140 | * | 0.0499 (0.017) | 0.0845 (0.034) |

| **FIB7** | *U* | *p* | sig. | **GULF** | HS |
|---|---|---|---|---|---|
| *I* | 84.0 | 0.5777 | ns | 0.305 (0.0338) | 0.288 (0.0804) |
| *Na* | 87.5 | 0.5376 | ns | 1.89 (0.0787) | 2.00 (0.000) |
| *uHe* | 88.0 | 0.6927 | ns | 0.181 (0.0238) | 0.172 (0.0620) |
| $H_O$ | 90.0 | 0.7538 | ns | 0.305 (0.0338) | 0.288 (0.0804) |
| $F_{IS}$ | 48.5 | 0.0949 | ns | 0.00904 (0.0246) | -0.0764 (0.0226) |

**TABLE S4.** Kruskal-Wallis statistics comparing genetic diversity values between the four colonies inside the Gulf: HA = Hawar, Bahrain; BU = Butina, UAE; W.I. = World Islands, UAE; Siniya Island, UAE. Comparisons are based on mean values estimated with GENALEX for cytochrome oxidase I (COI; top rows) and β-fibrinogen intron 7 (FIB7; bottom rows) across 10 and 7 SNPs, respectively. *Na* = number of different haplotypes (COI) or alleles (FIB7), *I* = Shannon's information index; *uh* = unbiased diversity (COI), $F_{IS}$ = fixation index (FIB7); $H_O$ = observed heterozygosity (FIB7), *uHe* = unbiased expected heterozygosity (FIB7). For Kruskal-Wallis tests: *KW* = Kruskal-Wallis statistic; ns = not significant. Mean values and standard errors of the means (in parentheses) are listed for each diversity measure per colony (HA, BU, W.I. and SI)—note that these are also tabulated in Table 2 of the main text.

| **COI** | ***KW*** | ***p*** | **sig.** | **HA** | BU | W.I. | SI |
|---|---|---|---|---|---|---|---|
| *Na* | 0.520 | 0.9145 | ns | 1.30 (0.153) | 1.20 (0.133) | 1.20 (0.133) | 1.30 (0.153) |
| *I* | 0.416 | 0.9368 | ns | 0.101 (0.000038) | 0.078 (0.000006) | 0.083 (0.000009) | 0.069 (0.000005) |
| *uh* | 0.374 | 0.9456 | ns | 0.0609 (0.039) | 0.0487 (0.038) | 0.0533 (0.039) | 0.0368 (0.022) |

| **FIB7** | ***KW*** | ***p*** | **sig.** | **HA** | BU | W.I. | SI |
|---|---|---|---|---|---|---|---|
| *Na* | 0.514 | 0.9157 | ns | 1.86 (0.143) | 1.86 (0.143) | 2.00 (0.218) | 1.86 (0.143) |
| *I* | 2.731 | 0.4350 | ns | 0.309 (0.0668) | 0.309 (0.0685) | 0.254 (0.0779) | 0.347 (0.0678) |
| $F_{IS}$ | 7.744 | 0.0516 | ns | -0.0457 (0.0312) | 0.0868 (0.0485) | -0.0782 (0.0228) | 0.0732 (0.0560) |
| $H_O$ | 2.603 | 0.4570 | ns | 0.184 (0.0411) | 0.172 (0.0520) | 0.162 (0.0669) | 0.192 (0.0410) |
| *uHe* | 2.731 | 0.4350 | ns | 0.182 (0.0459) | 0.183 (0.0474) | 0.148 (0.0587) | 0.211 (0.0459) |
